# CREP: Cis-Regulatory Element Predictor Based on Fine-Tuned Enformer

**DOI:** 10.64898/2026.06.05.730309

**Authors:** Nicolò Stranieri, Simone G. Riva, Jim R. Hughes

**Author notes:** These authors contributed equally to this work (Nicolò Stranieri, Simone G. Riva).

## Abstract

A substantial fraction of disease-associated genetic variants reside in non-coding regions of the genome, where they act by perturbing *cis*-regulatory elements (CREs) such as enhancers, promoters, and insulators. While recent sequence-based deep learning models, such as Enformer, accurately predict continuous epigenomic signals from DNA sequence, they do not directly provide discrete and interpretable CRE annotations. Here, we present CREP (*Cis*-Regulatory Element Predictor), a fine-tuned version of Enformer trained to predict regulatory element identity from sequence using REgulamentary-derived annotations across multiple human cell-types.

Through a controlled experimental framework, we show that incorporating diverse cell-types improves model performance. CREP leverages cell-type-specific training data to learn regulatory representations while producing a unified prediction of CRE identity from sequence. This is demonstrated by the Vanuatu SNP, a non-coding variant that creates a *de novo* erythroid regulatory element, which is correctly detected only when erythroid data are included during training.

Error analysis further reveals that apparent misclassifications between enhancers and promoters reflect their shared regulatory architecture, supporting the view of CREs as a functional continuum rather than strictly discrete classes. Together, these results demonstrate that CREP enables interpretable prediction of regulatory element identity from sequence and provides a framework for the functional interpretation of non-coding genetic variation.

## I. Introduction & Motivation

A substantial fraction of disease-associated genetic variants identified by genome-wide association studies (GWAS) reside in non-coding regions of the genome, where they are presumed to exert their effects by altering the activity of *cis*-regulatory elements (CREs) such as enhancers, promoters, and insulator elements [1]. These regulatory sequences play a central role in controlling gene expression by coordinating the recruitment of transcription factors (TFs) and chromatin-modifying complexes that determine when and where genes are transcribed. Unlike protein-coding mutations, which often directly disrupt gene function, non-coding variants frequently act by perturbing regulatory activity, for example, by altering TF binding sites or modifying chromatin accessibility. As a consequence, identifying which genomic regions function as active CREs, and in which cellular contexts they operate, is critical for understanding the mechanisms underlying both normal development and complex disease. The activity of CREs is highly dependent on cellular context. While promoters and CTCF boundary elements, characterised by the binding of the CCCTC-binding factor, tend to show relatively stable activity across related cell-types, enhancers are often strongly cell-type-specific and may only become active at particular developmental stages or under specific physiological conditions. This context dependence means that the regulatory potential encoded in DNA sequence cannot be interpreted independently of the cellular environment. In practice, CRE annotations are therefore derived from combinations of epigenomic assays, such as chromatin accessibility, histone modifications, and TF binding, that collectively indicate regulatory activity in specific cell-types. However, the resulting datasets are typically heterogeneous, assay-specific, and difficult to integrate into a unified catalogue of CREs suitable for computational modelling [2]–[5].

Sequence-based deep learning models have recently emerged as a powerful approach to learning the regulatory code directly from DNA. By training on large collections of functional genomics data, these models learn complex sequence patterns associated with regulatory activity and can predict genome-wide regulatory signals from sequence alone. Early approaches such as DeepSEA [6] and Basenji [7] demonstrated that convolutional neural networks can capture regulatory sequence features and predict a variety of functional genomic signals, including chromatin accessibility and TF binding.

More recent work has begun to extend these approaches by incorporating richer architectures and additional modalities. For example, transformer-based and multimodal frameworks have been proposed to jointly model DNA sequence and epigenomic signals, improving the ability to capture regulatory grammar and generalise across cellular contexts [8], [9]. These approaches highlight the potential of integrating sequence and chromatin information, but they typically remain focused on predicting functional signals or accessibility profiles rather than explicit CRE identities.

Building on these foundations, Enformer [10], a transformer-based architecture with a 196 kb receptive field, represents the current state of the art in this domain. Trained on more than 6,000 genomic assay tracks, Enformer predicts a wide range of continuous epigenomic and transcriptomic signals, including DNase accessibility, TF binding, and transcription initiation measured by CAGE. By modelling long-range genomic context, Enformer is able to capture distal regulatory interactions and learn sequence features that influence gene regulation across large genomic distances. Recent extensions such as REnformer [11] further refine this framework by explicitly incorporating cell-type-specific regulatory signals during training, enabling improved modelling of context-dependent regulatory activity while preserving long-range sequence representations. More recent models, such as AlphaGenome [12], further scale sequence-based prediction by integrating larger training corpora and improved architectural designs, achieving enhanced performance across diverse regulatory prediction tasks. Together, these models reflect a trend toward increasingly comprehensive modelling of genome regulation directly from sequence, with the ability to jointly predict multiple functional genomic modalities.

However, despite these advances, these approaches are fundamentally designed to predict continuous functional signals rather than discrete CRE identities. As a result, their outputs require downstream processing or heuristic thresholding to infer whether a given genomic region functions as an active enhancer, promoter, or CTCF element in a specific cellular context. While these models learn rich representations of regulatory sequence features, translating their outputs into biologically interpretable CRE identities remains a non-trivial step.

A further challenge lies in the lack of consistent training datasets for CREs prediction. Unlike genes, which have well-established annotations, CREs lack a universally agreed ground-truth catalogue. Existing resources are typically derived from individual assays or aggregated across multiple cell-types, making them difficult to use for training models that aim to predict cell-type-specific regulatory activity. To address this limitation, we leverage REgulamentary [13], a framework that integrates multiple chromatin features to generate cell-type-specific annotations of CREs. By combining information from chromatin accessibility, histone modifications, and TF binding profiles, REgulamentary produces unified binary annotations of promoters, enhancers, and CTCF-associated elements for individual cell-types, providing a consistent set of labels suitable for supervised learning.

Here we present CREP (*Cis*-Regulatory Element Predictor), a fine-tuned version of Enformer trained on high-quality REgulamentary annotations from multiple human cell types to predict unified CRE identity from sequence [13]. By reusing Enformer’s pre-trained long-range sequence representations and adapting them through fine-tuning to a binary classification objective, CREP inherits the contextual understanding of its parent model while learning to produce discrete and interpretable regulatory predictions. CREP is trained on cell-type-specific regulatory annotations but produces a unified prediction of CRE identity from sequence. An overview of the framework is presented in Figure 1.

**Figure 1.**
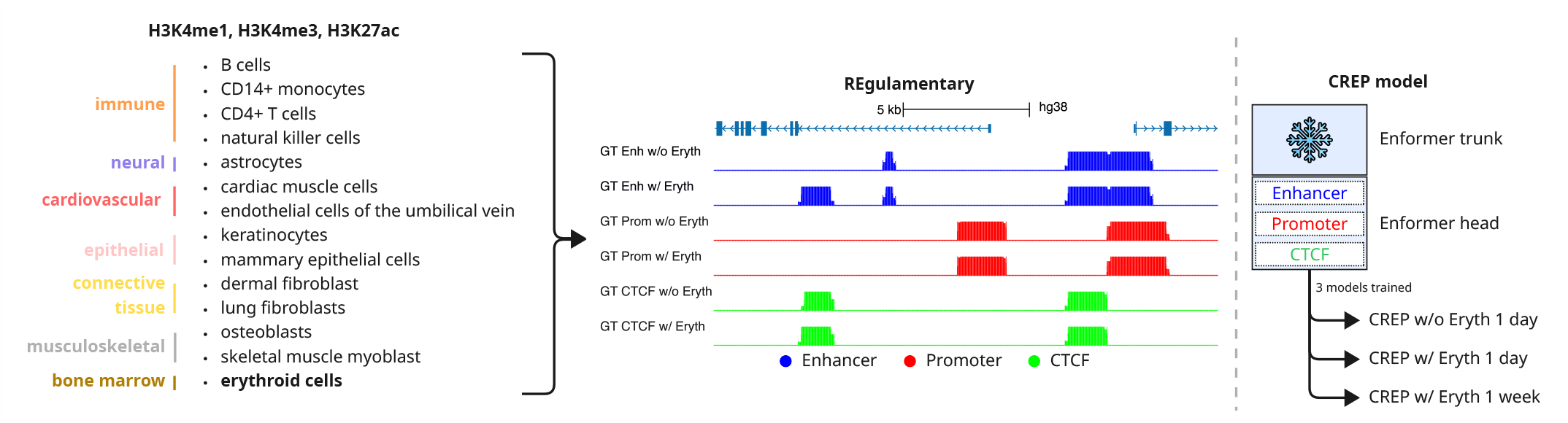
Overview of the CREP framework. *Left:* Cell-types used for training, spanning 13 human lineages, with erythroid cells added as a 14^th^ cell-type in a separate experiment. REgulamentary integrates H3K4me1, H3K4me3, and H3K27ac histone marks to generate genome-wide binary annotations of enhancers (blue), promoters (red), and CTCF sites (green). *Centre:* Example genomic locus showing ground-truth (GT) CRE annotations with and without erythroid data, illustrating cell-type-specific differences in regulatory element activity. *Right:* The CREP architecture. A frozen Enformer trunk provides long-range sequence representations, which are fed into a trainable Enformer head producing three output tracks (enhancer, promoter, CTCF). Three models are trained under different conditions: without erythroid data (1 day), with erythroid data (1 day), and with erythroid data (1 week).

We evaluate CREP by systematically varying training data composition and duration, culminating in a biological validation using the Vanuatu single nucleotide polymorphism (SNP), a well-characterised variant that creates a *de novo* erythroid regulatory element in the *α*-globin *locus* [14], [15].

## II. Data & Method

This section describes the data sources, annotation strategies, preprocessing pipeline, and model architecture used to train CREP.

### A. Data collection

CRE activity annotations were derived from publicly available chromatin and epigenomic datasets mapped to the human reference genome (hg38). Given the absence of a universally accepted ground-truth catalogue of CREs, we employed REgulamentary [13], a rule-based framework for *de novo* genome-wide annotation of CREs in a cell-type-specific manner. REgulamentary integrates multiple layers of epigenomic information, including chromatin accessibility, histone modification marks (H3K4me1, H3K4me3, H3K27ac), and CTCF ChIP-seq binding profiles. Based on a hierarchical decision process that combines these signals, each open chromatin region is assigned one of five regulatory identities: enhancer, promoter, CTCF, enhancer/CTCF, or promoter/CTCF. The hybrid classes (enhancer/CTCF and promoter/CTCF) capture regions exhibiting both regulatory activity and insulator-like properties, reflecting the complexity of chromatin organisation.

REgulamentary annotations were generated for 13 human cell-types spanning diverse lineages, including immune (B cells, CD14^+^ monocytes, CD4^+^ T cells, natural killer cells), neural (astrocytes), cardiovascular (cardiac muscle cells, endothelial cells of the umbilical vein), epithelial (keratinocytes, mammary epithelial cells), connective tissue (dermal and lung fibroblasts), and musculoskeletal (osteoblasts, skeletal muscle myoblasts) cell-types. This diversity enables the evaluation of cell-type-specific regulatory predictions across a broad biological spectrum. In a subsequent experiment, erythroid cell annotations were incorporated to assess the impact of including highly specialised, lineage-specific data on variant effect prediction.

### B. Data processing

For each annotated CRE, a genomic sequence window of 196 kb centred on the element midpoint was extracted, matching the input resolution of Enformer. Sequences were one-hot encoded as an *L×* 4 binary matrix representing the four nucleotides (A, C, G, T), where *L* = 196, 608 base pairs. This encoding preserves the raw nucleotide information while allowing efficient processing by convolutional and transformer-based architectures.

Regression targets were constructed directly from REgulamentary binary annotations. For each of the three CRE classes (enhancer, promoter, CTCF), a genome-wide binary annotation track was generated by assigning a value of 1 to each base pair overlapping an annotated element and 0 elsewhere. The output of CREP is defined over non-overlapping 128 bp bins, matching the resolution of the Enformer backbone. The regression target for each bin is therefore the sum of the binary annotation over the 128 constituent base pairs, yielding integer values in the range [0, 128]. A bin fully covered by a CRE of a given class receives a target value of 128, while a bin with no annotated overlap receives a value of 0. Bins at element boundaries receive intermediate values proportional to the overlap. This formulation naturally encodes both element presence and sub-bin boundary information, and is consistent with the output space of the original Enformer training objective.

To ensure a rigorous evaluation and prevent information leakage, genomic sequences were partitioned into non-overlapping training (80%), validation (10%), and test (10%) sets using the Basenji data preprocessing framework [16]. The genome was first divided into contiguous segments after excluding assembly gaps and blacklist regions (provided via the `-g` parameter). These segments were then randomly allocated to the three sets based on the specified proportions (`-t .1 -v .1`), ensuring that sequences in the test set are not trivially similar to those seen during training. The validation set was used for monitoring training progress and model selection, while all final performance metrics are reported exclusively on the held-out test set.

### C. Model architecture and training

CREP is implemented as a fine-tuned version of the pre-trained Enformer model. Rather than training a sequence model from scratch, we leverage Enformer’s learned representation of long-range genomic context and adapt it to a supervised prediction task. Specifically, the original Enformer output head, a linear projection to 5,313 human and 1,643 mouse signal tracks, was replaced with a task-specific regression head producing three output tracks of length 896, corresponding to the three CRE classes: enhancer, promoter, and CTCF.

Given an input sequence, the model produces a latent representation that captures both local sequence features and distal regulatory interactions. This representation is then mapped through the new output head to generate class-specific predictions across the genomic window, where each position corresponds to a 128 bp bin. The model is trained to predict, for each bin and each class, the number of base pairs covered by intervals of the corresponding CRE class.

Fine-tuning was performed using supervised learning with Huber loss as the regression objective [17]. All models were trained with a batch size of 16. To ensure deterministic and consistent embeddings across all experiments, the pre-trained Enformer backbone (excluding the prediction head) was set to evaluation mode with gradients disabled (`.eval()` and `no_grad()`), effectively freezing the learned representations. Only the task-specific output head was trained, allowing the model to leverage Enformer’s pre-trained genomic representations while adapting to the CRE prediction task. These hyperparameters and architectural choices were held constant across all experiments to isolate the effects of training data composition.

### D. Experimental design

Rather than focusing primarily on hyperparameter optimisation, we evaluate CREP through a controlled experimental ladder in which training data composition is systematically varied. Specifically, we compare models trained on 13 cell-types versus 14 cell-types (the latter including erythroid data), and models trained for 1 day versus 1 week.

The ladder culminates in a biological validation using the Vanuatu SNP, a well-characterised variant that creates a *de novo* erythroid regulatory element in the *α*-globin *locus* [14], [15].

## III. Results

We trained three CREP models, each initialised from the same pre-trained Enformer weights and fine-tuned on REgulamentary-derived annotations, systematically varying cell-type diversity (13 vs 14 cell-types) and training duration (1 day vs 1 week).

### A. Effect of cell-type diversity on model performance

To assess the impact of increasing biological diversity in the training data, we compared a baseline model trained on REgulamentary annotations from 13 cell-types for 1 day with a model including an additional erythroid cell-type (14 cell-types total). Erythroid cells represent a highly specialised lineage with distinct regulatory programs.

As shown in Figure 2a, the inclusion of erythroid data leads to measurable differences in model predictions after a single day of training. Furthermore, extending training duration from 1 day to 1 week (Figure 2b) allows us to assess whether these effects persist or are amplified with additional optimisation.

**Figure 2.**
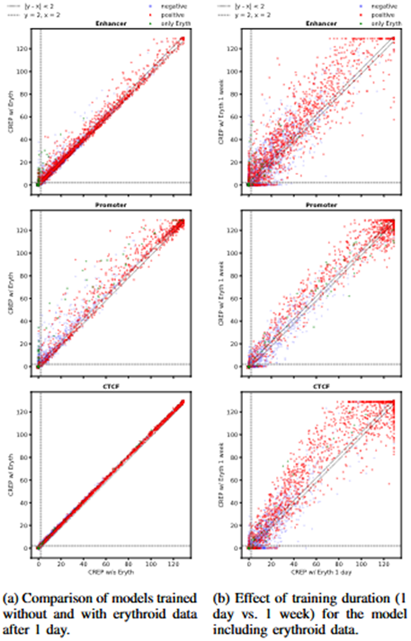
Impact of cell-type diversity and training duration on CREP performance. Predictions *>*129 bp are clipped to 129 for visualisation (see text). Points are coloured by ground truth: positive bins (red), negative bins (blue), and erythroid-specific bins (green). The diagonal dotted line indicates |*y* ™ *x*| *<* 2 bp agreement.

Because each regression target is defined as the count of annotated base pairs within a 128 bp bin (see Section II-B), the maximum attainable ground-truth value is 128. Predictions exceeding this ceiling, therefore, indicate overconfidence: the model assigns a value that is physically impossible given a fully covered bin. Such overconfident predictions are clipped to 129 for display purposes, mapping them to a uniform ceiling that does not distort the visual comparison between models while preserving all meaningful variation within the range [0, 129].

In both comparisons, the model with additional data or training time shows increased prediction confidence, with points predominantly above the diagonal (y-axis *>* x-axis). Notably, Figure 2a reveals that regions predicted as zero by the model without erythroid data (bottom-left quadrant) are assigned positive predictions by the erythroid-inclusive model, and these predictions correspond almost exclusively to erythroid-specific regions (green points). This demonstrates that the model successfully learns cell-type-specific regulatory patterns when provided with appropriate training data. In contrast, CTCF predictions remain largely unchanged between the two models, indicating that this class is already well-captured by the 13-cell-type baseline and is not substantially affected by the addition of erythroid data.

To quantify these effects, we evaluated model performance at two complementary levels: bin-level (128 bp resolution) and region-level (contiguous CRE elements). Bin-level metrics assess pixel-perfect prediction accuracy at the finest granularity, while region-level metrics evaluate whether the model successfully identifies the location of regulatory elements even when bin-level boundaries are imperfect. Figure 3 compares performance metrics between the 1 day and 1 week models, both trained with erythroid data.

**Figure 3.**
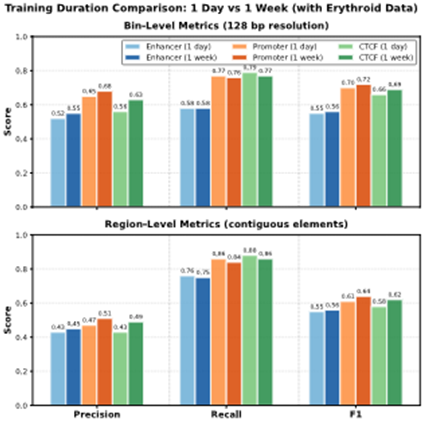
Comparison of classification metrics for models trained with erythroid data for 1 day (lighter bars) versus 1 week (darker bars). Top panel: bin-level metrics (128 bp resolution). Bottom panel: region-level metrics (contiguous elements). Extended training improves precision across both evaluation levels while maintaining high recall.

Extended training improves bin-level precision across all classes (enhancer: 0.52 *→* 0.55, promoter: 0.65 *→* 0.68, CTCF: 0.56 *→* 0.63) while maintaining similar recall, resulting in higher F1 scores. At the region level, precision increases from 0.43–0.47 to 0.45–0.51 across classes, while recall remains high and stable (0.75–0.88). This pattern reveals an important characteristic of the training dynamics: region-level recall is substantially higher than bin-level recall (0.76–0.88 vs 0.58–0.79), indicating that CREP successfully identifies the location of regulatory elements even when bin-level boundaries are imperfect. This highlights that region-level evaluation better reflects biological relevance, as regulatory elements are inherently contiguous and boundary definitions are approximate. Conversely, region-level precision is lower than bin-level precision (0.43–0.51 vs 0.52–0.68), reflecting the fact that false positive predictions grouped into discrete regions represent a larger fraction of total predicted regions than of total predicted bins. This highlights that region-level evaluation better reflects biological relevance, as regulatory elements are inherently contiguous and boundary definitions are approximate.

The improvement in precision without recall degradation indicates that extended training primarily refines prediction boundaries and reduces false positive rates rather than discovering additional regulatory elements. The model learns to be more selective in its predictions, consistent with better calibration of prediction confidence over time. Recall values remain consistently high across training durations, demonstrating that the model identifies the majority of regulatory elements early in training and that extended optimisation primarily refines prediction quality rather than expanding coverage.

### B. Effect of training duration on model convergence and performance

We compared the 14-cell-type model (including erythroid cells) trained for 1 day versus 1 week to evaluate whether CREP benefits from extended optimisation. All models were trained on a single NVIDIA L40S GPU.

Figure 4 shows training and validation curves for both loss and R^2^ over the course of training. The vertical line at 1 day marks the point at which the shorter training run was stopped. Both loss and R^2^ show steady improvement over the first day of training, with validation metrics closely tracking training metrics, indicating no overfitting. Loss decreases from approximately 2.6 to 1.8, while R^2^ increases from near zero to approximately 0.45 on the validation set. Beyond the 1day mark, training continues to improve model performance, with validation loss decreasing further and R^2^ continuing to increase, eventually plateauing near 0.45. The close alignment between training and validation curves throughout the entire training period demonstrates that the model generalises well to held-out data and that extended training refines predictions without overfitting to the training set.

**Figure 4.**
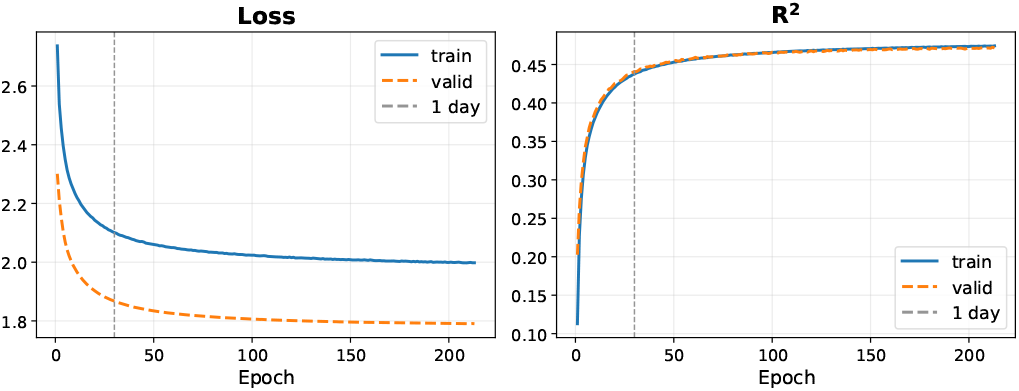
Training and validation curves for the 14-cell-type model. Loss (left) and R^2^ (right) are shown over 200 epochs. The vertical line indicates 1 day of training. Both metrics show continued improvement beyond 1 day, with validation performance closely tracking training performance, indicating effective generalisation without overfitting.

### C. Error analysis

To better understand the behaviour of CREP predictions, we performed a detailed analysis of prediction outcomes on the held-out test set across the three CRE classes: enhancer, promoter, and CTCF. Rather than focusing solely on errors, this analysis reveals how the model captures the underlying regulatory landscape and where discrepancies with the reference annotations arise.

As shown in Figure 5, prediction patterns differ across CRE classes in ways that reflect both biological properties and annotation conventions. CTCF sites show the most consistent behaviour, with a high number of true positives and comparatively fewer ambiguous predictions. This is expected, as CTCF binding is strongly driven by a well-defined sequence motif and is associated with relatively stable and structurally constrained genomic functions, making it more readily predictable from sequence alone.

**Figure 5.**
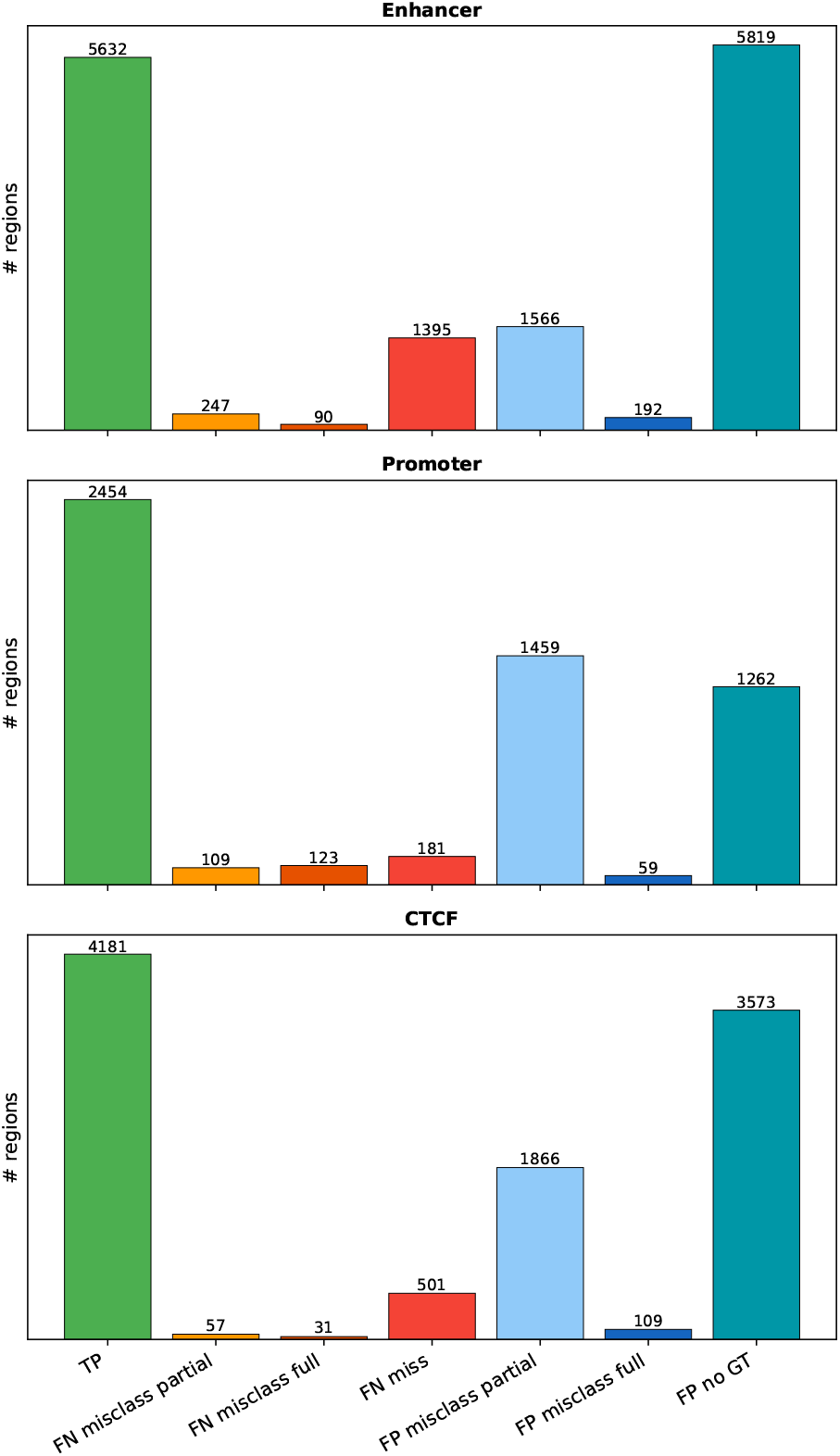
Breakdown of prediction outcomes across CRE classes. Categories include true positives (TP), partial and full false negative misclassifications, missed elements (FN miss), and false positives, including partial misclassification, full misclassification, and predictions without corresponding annotated ground truth (FP no GT).

In contrast, enhancer and promoter predictions exhibit a higher degree of apparent misclassification. However, these discrepancies should be interpreted in light of the well-established functional and architectural overlap between these two classes. Promoters and enhancers share many molecular features, including TF binding, chromatin accessibility, and the presence of activating histone marks such as H3K27ac. Moreover, bidirectional transcription and enhancer RNA (eRNA) production have been observed at active enhancers, blurring the distinction between enhancers and promoters [18]–[20]. Conversely, promoters can exhibit enhancer-like activity in certain contexts, acting as distal CREs for other genes [21], [22].

This ambiguity is further illustrated in Figure 6, which focuses on full misclassification events. Rather than being randomly distributed, these errors follow a structured pattern. In particular, enhancer regions are frequently predicted as promoters, and promoter regions are often predicted as enhancers, indicating a bidirectional confusion between these two classes. In contrast, confusion involving CTCF is more limited and asymmetric, with fewer promoter-to-CTCF or enhancer-to-CTCF misassignments. This suggests that the model has learned a coherent representation of CRE space, in which enhancers and promoters occupy a closely related functional continuum, while CTCF sites form a more distinct category.

**Figure 6.**
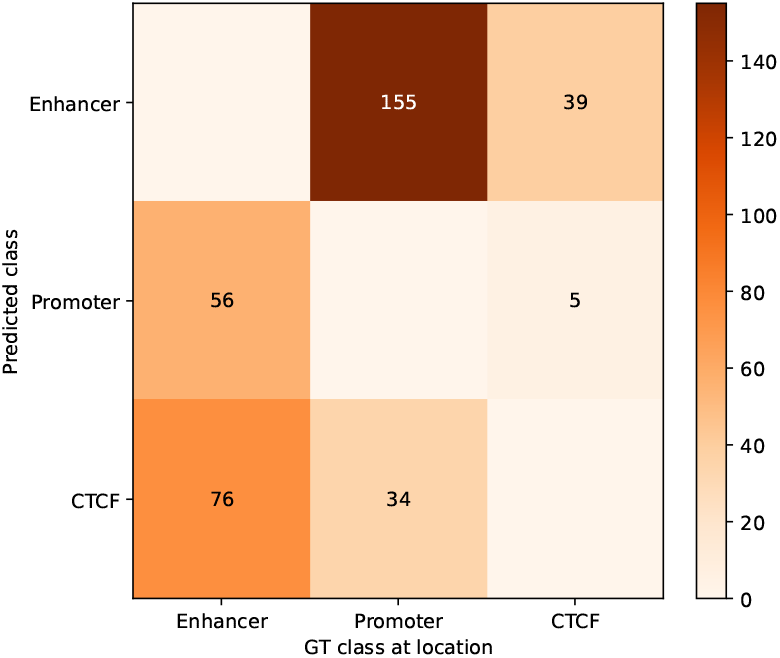
Confusion matrix for full misclassification events, showing the relationship between predicted and ground-truth CRE classes.

As a result, regions classified as misclassified between enhancer and promoter categories may reflect genuine biological ambiguity rather than model error. Differences in annotation are also influenced by the operational definitions used in peak-calling pipelines [23], [24] and labelling strategies, which may assign regions to one class or the other based on proximity to transcription start sites or specific combinations of epigenomic marks.

We also observe a substantial number of predictions that do not overlap annotated CREs (FP no GT). Importantly, these predictions should not be interpreted as spurious. The reference annotations used in this study are derived from peak-calling approaches applied to experimental data, which are inherently incomplete and dependent on assay sensitivity and thresholding. Sequence-based models such as CREP are not constrained by these limitations and may identify regions with regulatory potential that are not captured in the available datasets, including regions with weak, transient, or context-specific activity.

Across all classes, the balance between matched predictions, class ambiguity, and predictions without annotated support highlights both the strengths of CREP and the limitations of current CRE annotation frameworks. Overall, these results demonstrate that the model effectively captures core regulatory signals, particularly for well-defined elements such as CTCF, while also reflecting the inherent continuum between enhancer and promoter activity in the human genome.

### D. Variant effect prediction: the Vanuatu SNP

A key motivation for CRE prediction is the functional interpretation of non-coding genetic variants. The Vanuatu SNP is a T-to-C single nucleotide change in the *α*-globin *locus*, first identified in individuals from Melanesia, which creates a novel GATA-1 binding site and establishes a *de novo* CRE active specifically in erythroid cells, leading to *α*-thalassaemia [14], [15].

Variant effect prediction was performed *in silico* by running CREP on the 196 kb genomic window centred on the Vanuatu *locus* with either the reference (T) or alternate (C) allele, and computing the change in predicted CRE signal (ISM Δ-score) at the relevant genomic bin. We evaluated both RE models trained with and without erythroid data.

Without erythroid training data, CREP produced no meaningful difference in predicted regulatory activity between the reference and alternate alleles, suggesting that the model lacks the erythroid-specific regulatory grammar required to recognise the newly introduced GATA-1 motif as functionally relevant. In contrast, after incorporating erythroid annotations during training, CREP predicts a clear gain of regulatory activity at the Vanuatu *locus* specifically in erythroid cells, with no comparable signal observed in other cell-types. This demonstrates that accurate variant effect prediction critically depends on the inclusion of appropriate cellular context. This result highlights that regulatory grammar is inherently cell-type-dependent and must be learned from appropriately matched training data.

Notably, CREP classifies the gained regulatory activity as enhancer-like rather than promoter-like. While this appears to differ from the original characterisation of the *locus*, this distinction is not contradictory. The boundary between enhancers and promoters is increasingly recognised as fluid, with both classes sharing common molecular features such as TF binding, chromatin accessibility, and activating histone marks [18], [19]. Moreover, CREs can exhibit dual functionality, with promoters acting as enhancers in certain contexts and enhancers displaying promoter-like transcriptional activity [21], [22]. From this perspective, the prediction of enhancer activity is consistent with the emergence of a functional CRE driven by the newly created TF binding site.

This result highlights that correct functional interpretation of regulatory variants does not require strict agreement on enhancer versus promoter labels, but rather accurate identification of context-specific regulatory activity, consistent with the view of CREs as a functional continuum [19].

To further interpret the model prediction, we analysed both the predicted CRE tracks and the underlying sequence contributions using feature attribution with DeepLIFT [25], [26]. As shown in Figure 7, the genome browser view reveals a clear gain of regulatory activity at the Vanuatu *locus* in the alternate allele, specifically in models trained with erythroid data, while no comparable signal is observed without erythroid training. This effect is consistent across training durations and confined to the local genomic context of the variant.

**Figure 7.**
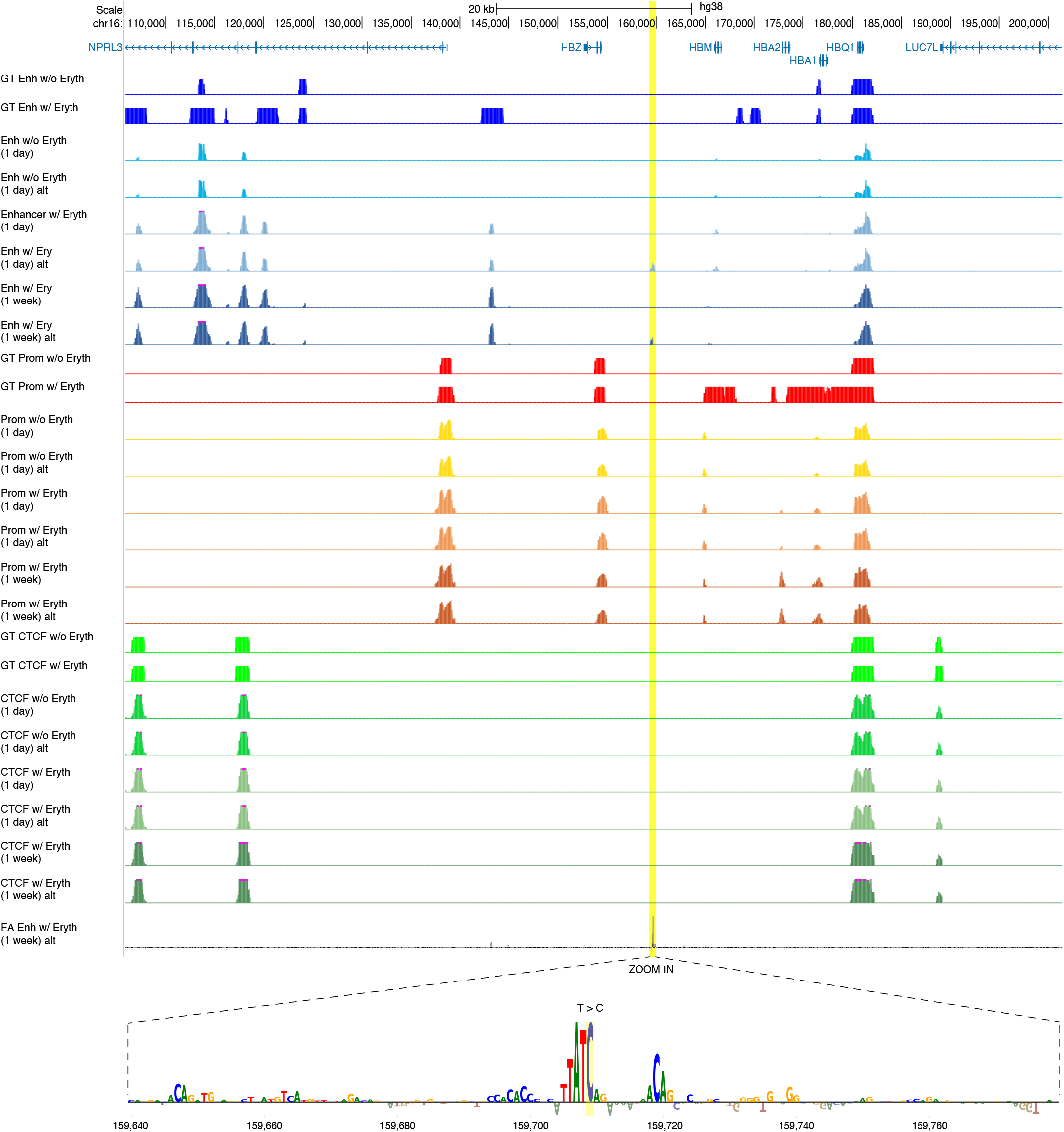
Variant effect prediction at the Vanuatu *locus*. Genome browser view across ∼ 100 kb window showing ground-truth annotations and CREP predictions for enhancer, promoter, and CTCF tracks under different training conditions (without erythroid data, with erythroid data after 1 day, and after 1 week). Enhancer, promoter, and CTCF signals are shown in shades of blue, red, and green, respectively. For each configuration, predictions are displayed for both reference (T) and alternate (C) alleles. The Vanuatu SNP position is highlighted (yellow), and a gain of regulatory activity is observed specifically in the erythroid-trained models for the alternate allele, demonstrating that detection of the regulatory effect depends on the inclusion of erythroid-specific training data. Bottom: zoom-in of the variant *locus* with DeepLIFT feature attribution, showing a strong contribution centred on the mutation and consistent with the creation of a GATA binding motif.

The zoomed attribution profile further shows that the signal is strongly localised around the variant position and highlights the emergence of a GATA binding motif in the alternate allele. Together, these results provide mechanistic support for the prediction, indicating that CREP has learned to associate GATA motif instances with erythroid regulatory activity and to translate this sequence change into a context-dependent gain of regulatory function, effectively mapping sequence-level changes to functional regulatory outcomes in a cell-type-dependent manner.

Taken together, these results demonstrate that CREP not only detects the functional impact of a non-coding variant but also captures the underlying sequence determinants driving regulatory activity. Importantly, the model’s prediction aligns with biological expectations even when strict categorical labels (enhancer vs promoter) are ambiguous, reinforcing the view that regulatory element activity is better represented as a continuum rather than as discrete classes.

## IV. Conclusion & Future Work

In this work, we introduced CREP, a fine-tuned version of Enformer designed to predict CRE identity from DNA sequence, trained on annotations derived from multiple cell types. By leveraging REgulamentary-derived binary annotations, CREP bridges the gap between sequence-based prediction of functional signals and the explicit identification of CREs such as enhancers, promoters, and CTCF sites from sequence using cell-type-resolved training annotations.

We demonstrated that cell-type diversity and training duration both influence model performance. CREP generalises regulatory representations across cell-types while benefiting from cell-type-specific training data when available. This is most clearly illustrated by the Vanuatu SNP, where correct prediction of the regulatory effect was only achieved after incorporating erythroid-specific training data. This demonstrates that accurate modelling of regulatory variants requires learning cell-type-specific regulatory grammar directly from sequence.

Our analysis also highlights important limitations. First, the number of available training cell-types remains relatively limited, which constrains the diversity of regulatory programs that the model can learn. Second, the use of discrete binary labels does not fully capture the graded and context-dependent nature of regulatory activity, particularly for enhancers. Third, although CREP leverages a 196 kb receptive field inherited from Enformer, regulatory interactions can extend beyond this window and involve higher-order chromatin organisation that is not explicitly modelled.

These limitations point to several promising directions for future work. Scaling the approach to a larger and more diverse set of cell-types will be essential to improve generalisation and enable broader applicability. Extending the model to a multi-task setting, where CRE identity prediction is jointly learned with continuous regulatory signals, may provide a more faithful representation of regulatory activity. Incorporating three-dimensional genome organisation, for example through Hi-C [27] or related chromatin conformation data, could further improve the modelling of long-range regulatory interactions. Finally, applying CREP to the systematic prioritisation of non-coding variants identified in GWAS represents a key avenue for translating these models into practical tools for human genetics and disease research.

## Acknowledgment

N.S. is supported by the Wellcome Trust grants (225220/Z/22/Z). S.G.R. is supported by the MRC grant (MCUU00029/3). J.R.H. is supported by the Wellcome Trust grants (225220/Z/22/Z and 106130/Z/14/Z) and the MRC grant (MCUU00029/3).

## Declaration

J.R.H. is a co-founder and director of Nucleome Therapeutics and provides consultancy to the company.

## References

[1] M. T. Maurano, R. Humbert, E. Rynes, R. E. Thurman, E. Haugen, H. Wang, A. P. Reynolds, R. Sandstrom, H. Qu, J. Brody et al., “Systematic localization of common disease-associated variation in regulatory dna,” Science, vol. 337, no. 6099, pp. 1190–1195, 2012.

[2] D. J. Downes and J. R. Hughes, “Natural and experimental rewiring of gene regulatory regions,” Annual Review of Genomics and Human Genetics, vol. 23, pp. 73–97, 2022.

[3] M. S. Larke, R. Schwessinger, T. Nojima, J. Telenius, R. A. Beagrie, D. J. Downes, A. M. Oudelaar, J. Truch, B. Graham, M. Bender et al., “Enhancers predominantly regulate gene expression during differentiation via transcription initiation,” Molecular cell, vol. 81, no. 5, pp. 983–997, 2021.

[4] A. M. Oudelaar and D. R. Higgs, “The relationship between genome structure and function,” Nature Reviews Genetics, vol. 22, no. 3, pp. 154–168, 2021.

[5] T. H. Kim, Z. K. Abdullaev, A. D. Smith, K. A. Ching, D. I. Loukinov, R. D. Green, M. Q. Zhang, V. V. Lobanenkov, and B. Ren, “Analysis of the vertebrate insulator protein ctcf-binding sites in the human genome,” Cell, vol. 128, no. 6, pp. 1231–1245, 2007.

[6] J. Zhou and O. G. Troyanskaya, “Predicting effects of noncoding variants with deep learning–based sequence model,” Nature methods, vol. 12, no. 10, pp. 931–934, 2015.

[7] L. Barbadilla-Martínez, N. Klaassen, B. van Steensel, and J. de Ridder, “Predicting gene expression from dna sequence using deep learning models,” Nature Reviews Genetics, vol. 26, no. 10, pp. 666–680, 2025.

[8] S. A. Khan, X. Martínez-de Morentin, A. R. Alsabbagh, A. Maillo, Lagani, D. Gomez-Cabrero, R. Lehmann, and J. Tegner, “Multimodal foundation transformer models for multiscale genomics,” Nature Methods, pp. 1–13, 2025.

[9] M. E. Consens, C. Dufault, M. Wainberg, D. Forster, M. Karimzadeh, H. Goodarzi, F. J. Theis, A. Moses, and B. Wang, “Transformers and genome language models,” Nature Machine Intelligence, vol. 7, no. 3, pp. 346–362, 2025.

[10] Ž. Avsec, V. Agarwal, D. Visentin, J. R. Ledsam, A. Grabska-Barwinska, K. R. Taylor, Y. Assael, J. Jumper, P. Kohli, and D. R. Kelley, “Effective gene expression prediction from sequence by integrating long-range interactions,” Nature methods, vol. 18, no. 10, pp. 1196–1203, 2021.

[11] S. G. Riva, E. Sanders, T. Wilson, N. Stranieri, E. R. Gür, M. Baxter, and J. R. Hughes, “Renformer, a single-cell atac-seq predicting model to investigate open chromatin sites,” in 2025 IEEE Conference on Computational Intelligence in Bioinformatics and Computational Biology (CIBCB). IEEE, 2025, pp. 1–8.

[12] Ž. Avsec, N. Latysheva, J. Cheng, G. Novati, K. R. Taylor, T. Ward C. Bycroft, L. Nicolaisen, E. Arvaniti, J. Pan et al., “Advancing regulatory variant effect prediction with alphagenome,” Nature, vol. 649, no. 8099, pp. 1206–1218, 2026.

[13] S. G. Riva, E. Sanders, E. Georgiades, S. S. Venkatesh, M. Sergeant, E. Ravza Gür, J. C. Herrmann, M. Baxter, and J. R. Hughes, “Deciphering cis-regulatory elements using regulamentary,” Bioinformatics Advances, p. vbag079, 03 2026. [Online]. Available: 10.1093/bioadv/vbag079

[14] M. De Gobbi, V. Viprakasit, J. R. Hughes, C. Fisher, V. J. Buckle, H. Ayyub, R. J. Gibbons, D. Vernimmen, Y. Yoshinaga, P. De Jong et al., “A regulatory snp causes a human genetic disease by creating a new transcriptional promoter,” Science, vol. 312, no. 5777, pp. 1215–1217, 2006.

[15] Y. K. Bozhilov, D. J. Downes, J. Telenius, A. Marieke Oudelaar, N. Olivier, J. C. Mountford, J. R. Hughes, R. J. Gibbons, and D. R. Higgs, “A gain-of-function single nucleotide variant creates a new promoter which acts as an orientation-dependent enhancer-blocker,” Nature communications, vol. 12, no. 1, p. 3806, 2021.

[16] D. R. Kelley, “Cross-species regulatory sequence activity prediction,” PLoS computational biology, vol. 16, no. 7, p. e1008050, 2020.

[17] P. J. Huber, “Robust Estimation of a Location Parameter,” The Annals of Mathematical Statistics, vol. 35, no. 1, pp. 73 –101, 1964. [Online]. Available: 10.1214/aoms/1177703732

[18] T.-K. Kim and R. Shiekhattar, “Architectural and functional commonalities between enhancers and promoters,” Cell, vol. 162, no. 5, pp. 948–959, 2015.

[19] R. Andersson and A. Sandelin, “Determinants of enhancer and promoter activities of regulatory elements,” Nature Reviews Genetics, vol. 21, no. 2, pp. 71–87, 2020.

[20] R. Andersson, C. Gebhard, I. Miguel-Escalada, I. Hoof, J. Bornholdt, M. Boyd, Y. Chen, X. Zhao, C. Schmidl, T. Suzuki et al., “An atlas of active enhancers across human cell types and tissues,” Nature, vol. 507, no. 7493, pp. 455–461, 2014.

[21] J. M. Engreitz, J. E. Haines, E. M. Perez, G. Munson, J. Chen, M. Kane, P. E. McDonel, M. Guttman, and E. S. Lander, “Local regulation of gene expression by lncrna promoters, transcription and splicing,” Nature, vol. 539, no. 7629, pp. 452–455, 2016.

[22] O. Mikhaylichenko, V. Bondarenko, D. Harnett, I. E. Schor, M. Males, R. R. Viales, and E. E. Furlong, “The degree of enhancer or promoter activity is reflected by the levels and directionality of erna transcription,” Genes & development, vol. 32, no. 1, pp. 42–57, 2018.

[23] Y. Zhang, T. Liu, C. A. Meyer, J. Eeckhoute, D. S. Johnson, B. E. Bernstein, C. Nusbaum, R. M. Myers, M. Brown, W. Li et al., “Model-based analysis of chip-seq (macs),” Genome biology, vol. 9, no. 9, p. R137, 2008.

[24] L. D. Hentges, M. J. Sergeant, C. B. Cole, D. J. Downes, J. R. Hughes, and S. Taylor, “LanceOtron: a deep learning peak caller for genome sequencing experiments,” Bioinformatics, 07 2022, btac525. [Online]. Available: 10.1093/bioinformatics/btac525

[25] A. Shrikumar, P. Greenside, and A. Kundaje, “Learning important features through propagating activation differences,” in International conference on machine learning. PMlR, 2017, pp. 3145–3153.

[26] N. Kokhlikyan, V. Miglani, M. Martin, E. Wang, B. Alsallakh, J. Reynolds, A. Melnikov, N. Kliushkina, C. Araya, S. Yan et al., “Cap-tum: A unified and generic model interpretability library for pytorch. arxiv,” arXiv preprint 2009.07896, vol. 2, p. 5, 2020.

[27] E. Lieberman-Aiden, N. L. Van Berkum, L. Williams, M. Imakaev, T. Ragoczy, A. Telling, I. Amit, B. R. Lajoie, P. J. Sabo, M. O. Dorschner et al., “Comprehensive mapping of long-range interactions reveals folding principles of the human genome,” science, vol. 326, no. 5950, pp. 289–293, 2009.

